# Vagus nerve stimulation (VNS) modulates synaptic plasticity in the rat infralimbic cortex via Trk-B receptor activation to reduce drug-seeking

**DOI:** 10.1101/2024.01.25.577293

**Authors:** Christopher M. Driskill, Jessica E. Childs, Aarron J. Phensy, Sierra R. Rodriguez, John T. O’Brien, Kathy L. Lindquist, Aurian Naderi, Bogdan Bordieanu, Jacqueline F. McGinty, Sven Kroener

## Abstract

Drugs of abuse cause changes in the prefrontal cortex (PFC) and associated regions that impair inhibitory control over drug-seeking. Breaking the contingencies between drug-associated cues and the delivery of the reward during extinction learning reduces relapse. Vagus nerve stimulation (VNS) has previously been shown to enhance extinction learning and reduce drug-seeking. Here we determined the effects of VNS-mediated release of brain-derived neurotrophic factor (BDNF) on extinction and cue-induced reinstatement in rats trained to self-administer cocaine. Pairing 10 days of extinction training with VNS facilitated extinction and reduced drug-seeking behavior during reinstatement. Rats that received a single extinction session with VNS showed elevated BDNF levels in the medial PFC as determined via an enzyme-linked immunosorbent assay (ELISA). Systemic blockade of Tropomyosin receptor kinase B (TrkB) receptors during extinction, via the TrkB antagonist ANA-12, decreased the effects of VNS on extinction and reinstatement. Whole-cell recordings in brain slices showed that cocaine self-administration induced alterations in the ratio of AMPA and NMDA receptor-mediated currents in layer 5 pyramidal neurons of the infralimbic cortex (IL). Pairing extinction with VNS reversed cocaine-induced changes in glutamatergic transmission by enhancing AMPAR currents, and this effect was blocked by ANA-12. Our study suggests that VNS consolidates extinction of drug-seeking behavior by reversing drug-induced changes in synaptic AMPA receptors in the IL, and this effect is abolished by blocking TrkB receptors during extinction, highlighting a potential mechanism for the therapeutic effects of VNS in addiction.

**Significance Statement:** Extinction training can reverse maladaptive neuroplasticity induced by drugs of abuse, but adjunct treatments are sought that can facilitate the process and consolidate the newly formed memories. Pairing extinction training with vagus nerve stimulation (VNS) facilitates extinction and reduces drug-seeking behavior during reinstatement. Here, we show that rats receiving a single extinction session with VNS exhibit elevated brain-derived neurotrophic factor (BDNF) levels in the medial prefrontal cortex (mPFC). We also demonstrate that VNS consolidates the extinction of drug-seeking behavior by reversing cocaine-induced changes in synaptic AMPA receptors in the infralimbic cortex (IL) of the mPFC. This effect is blocked by the TrkB antagonist ANA-12, emphasizing the role of BDNF and TrkB receptors in the therapeutic effects of VNS in addiction.

## Introduction

Exposure to drug-associated cues or stress induces craving and relapse in abstinent patients with substance use disorder (Ehrman et al., 1992). During extinction learning, new associations are formed that compete with these triggers to inhibit behavioral responses (Millan et al., 2011). However, extinction training alone is often insufficient to prevent relapse (Conklin and Tiffany, 2002; Weiss et al., 2001), potentially because the corticolimbic networks that regulate cue- and reward processing, which include areas in the medial prefrontal cortex (mPFC), the amygdala, and the ventral striatum, become dysregulated by drug use (Fowler et al., 2007; Liu et al., 2009; Nic Dhonnchadha and Kantak, 2011). Modulating extinction processes and consolidating the newly-formed memories is therefore clinically relevant to reshape maladaptive behavior and prevent relapse (Taylor et al., 2009).

Vagus nerve stimulation (VNS) is clinically used to treat epilepsy (Ben-Menachem et al., 1994) and depression (Rush et al., 2005), but is considered for the treatment of an expanding range of neurologic conditions (Dawson et al., 2021; Tyler et al., 2017). VNS causes an increase in the levels and/or release of several neuromodulators, including brain-derived neurotrophic factor (BDNF) (Dorr and Debonnel, 2006; Follesa et al., 2007; Manta et al., 2013; Nichols et al., 2011; Roosevelt et al., 2006), which modulate cortical plasticity (Cao et al., 2016; Collins et al., 2021), and this plasticity can facilitate extinction training and reduce cue-induced reinstatement of drug-seeking (Childs et al., 2017; Childs et al., 2019; Liu et al., 2011). However, the networks and neurotransmitters that support these effects are still poorly understood.

Cocaine self-administration induces aberrant plasticity at glutamatergic synapses in the mPFC, leading to altered salience attribution and cue-induced drug-seeking (Caffino et al., 2018; Otis and Mueller, 2017; Sasase et al., 2019). Repeated exposure to cocaine and withdrawal also lower the level of BDNF in the mPFC (Fumagalli et al., 2007; McGinty et al., 2010; Pitts et al., 2016). BDNF modulates cocaine-induced changes in glutamatergic transmission in the mPFC (Barry and McGinty, 2017; Berglind et al., 2007; Go et al., 2016; Otis et al., 2014; Otis and Mueller, 2017) and supplementation of BDNF via a single infusion into the mPFC directly after a final cocaine self-administration session reduces cue- and cocaine prime-induced reinstatement following extinction or abstinence (Barry and McGinty, 2017; Berglind et al., 2007; Berglind et al., 2009; Go et al., 2016; Whitfield et al., 2011). We hypothesized that cocaine self-administration alters glutamatergic signaling in the IL, an area in the mPFC that is important for the expression of extinction memories (Augur et al., 2016; Gutman et al., 2017; Muller Ewald et al., 2019; Peters et al., 2009). We also hypothesized that pairing extinction training with VNS leads to BDNF release in the mPFC which can reverse these changes and reduce reinstatement. Our data show that VNS-evoked release of BDNF consolidates extinction of drug-seeking behavior by reversing drug-induced changes in synaptic AMPA receptors in the IL, and this effect is abolished by blocking tropomyosin receptor kinase B (TrkB) receptors during extinction.

## Materials and Methods

### Subjects

We used male Sprague-Dawley rats (Taconic, Germantown, NY) that were at least 80 days old (250-300g) at the time of surgery. Rats were individually housed and kept on a 12-hr. reverse light/dark cycle, with free access to food and water until surgery, when food was restricted to 25 g/day standard rat chow. All protocols were approved by the IACUC of The University of Texas at Dallas and were conducted in compliance with the NIH Guide for the Care and Use of Laboratory Animals.

### Drug Self-Administration and Extinction Training

Drug self-administration and extinction training were performed as previously described (Childs et al., 2017; Childs et al., 2019): Male Sprague Dawley rats (>80 days) were anesthetized and implanted with a catheter in the right external jugular vein for drug administration. During the same surgery, a custom-made cuff electrode was placed around the left vagus nerve for the delivery of VNS (Childs et al., 2015). Five to seven days following surgery, rats were trained in a single overnight session to self-administer food pellets (45 mg, Bio Serv, Flemmington, NJ) in an operant conditioning chamber (Med Associates, Saint Albans, VT). Drug self-administration training took place in the same chamber, which was equipped with two levers, a house light, a cue light, and a tone. Each active lever press produced a 0.05 ml infusion of 2.0 mg/mL cocaine (Sigma, St. Louis, MO) in saline, and the presentation of drug-paired cues (illumination of the light over the active lever and the presentation of a 2900 Hz tone), followed by a 20 s timeout. Self-administration sessions ended after 2 hours. Both right and left levers were available for the duration of the session and drug-seeking behavior was quantified as active lever presses. Rats self-administered cocaine for 15-18 days, during which they had to achieve at least 20 infusions per session. Subjects in the extinction groups underwent 10 days of extinction training in which lever presses on the previously active lever no longer produced cocaine or presentation of drug-paired cues. During extinction training rats received either sham-stimulation or VNS. Non-contingent VNS stimulation occurred every 5 min for 30 sec at 0.4 mA for the duration of the training session. These stimulation parameters are similar to those previously used to enhance retention performance in rats (Clark et al., 1995) and humans (Clark et al., 1999). After 10 days of extinction training, drug-seeking behavior was reinstated by presentation of the drug-associated cues in the operant conditioning chambers. During the reinstatement session presses on the “active” lever led to presentation of the previously drug-associated tone and light, but did not result in drug delivery or VNS. Additional control groups of rats receive yoked saline injections during the self-administration sessions, followed by sham- or VNS-treatment during the extinction period, to control for the pharmacological effects of cocaine and repeated VNS, respectively.

#### TrkB receptor antagonism

ANA-12 systemic intraperitoneal injections of the TrkB receptor antagonist ANA-12 (0.5 mg/kg in DMSO, or DMSO-vehicle at a volume of 0.5 ml/kg) during the 10 days of extinction learning (but not during the reinstatement session)

### Electrophysiology

Rats were anesthetized with an overdose of urethane (3 g/kg body weight; Fisher Scientific) and transcardially perfused for one minute with ice-cold oxygenated (95% O2, 5% CO2) cutting ACSF, consisting of (in mM): 110 choline (Sigma-Aldrich), 25 NaHCO3 (Fisher Scientific), 1.25 NaH2PO4 (Fisher Scientific), 2.5 KCl (Sigma-Aldrich), 7 MgCl2 (Sigma-Aldrich), 0.5 CaCl2 (Sigma-Aldrich), 10 dextrose (Fisher Scientific), 1.3 L-ascorbic acid (Fisher Scientific), and 2.4 Na+-pyruvate (Sigma-Aldrich). Brains were extracted and coronal sections (350 um) of the frontal cortex were cut on a vibratome (VT1000S, Leica) in cutting ACSF. Slices were transferred to a holding chamber containing warmed (35C) recording ACSF and cooled to room temperature over a one-hour period. The recording ACSF consisted of (in mM): 126 NaCl (Fisher Scientific), 25 NaHCO3, 1.25 NaH2PO4, 2.5 KCl, 2 MgCl2, 2 CaCl2, 10 dextrose, 2.4 Na+-pyruvate, and 1.3 L-ascorbic acid. For data collection, slices were transferred to a recording chamber affixed to an Olympus BX61WI microscope (Olympus) with continuous perfusion of oxygenated recording ACSF at room temperature. Whole-cell voltage-clamp recordings were obtained from pyramidal cells in the IL using Axon Multiclamp 700B amplifiers (Molecular Devices). Data were acquired and analyzed using AxoGraph X (AxoGraph Scientific). Recording electrodes (WPI; 4–6 MΩ open tip resistance) were filled with an internal solution consisting of (in mM): 130 CsCl (Sigma-Aldrich), 20 tetraethylammonium chloride (Sigma-Aldrich), 10 HEPES (Sigma-Aldrich), 2 MgCl2, 0.5 EGTA (Sigma-Aldrich), 4 Mg2+-ATP (Sigma-Aldrich), 0.3 Lithium-GTP (Sigma-Aldrich), 14 phosphocreatine (Sigma-Aldrich), and 2 QX-314 bromide (Tocris Bioscience). Theta-glass pipettes (Warner Instruments) connected to a stimulus isolator (WPI) were used for focal stimulation of synaptic potentials. Access resistance was monitored throughout the recording, and a <20% change was deemed acceptable. EPSCs were isolated by blocking chloride channels with the addition of picrotoxin (75uM; Sigma-Aldrich) into the recording ACSF. The ratio of currents through AMPA or NMDA receptors (AMPAR:NMDAR) was obtained by clamping cells at +40mv holding potential and applying local electrical stimulation. A compound evoked EPSC (eEPSC) was first recorded, then the AMPA component was isolated by washing CPP ((+/-)-3-(2-carboxypiperazin-4-yl)propyl-1-phosphonic acid; 10 uM; Sigma-Aldrich) into the bath. A minimum of 20 sweeps each were average for the compound and AMPA-only eEPSCs. The NMDA component was then obtained by digital subtraction of the AMPA component from the compound trace. The peak amplitude of the NMDA and AMPA traces were used to calculate the NMDAR/AMPAR ratio. For all recordings a minimum of 4 rats were used per treatment group.

### BDNF ELISA

One cohort of rats was killed immediately after a single day of extinction training with VNS or sham-stimulation to determine VNS-induced levels of BDNF in the mPFC via an enzyme-linked immunosorbent assay (ELISA). Brains were extracted and frozen in 2,3-Methylbutane on dry ice and stored at −80C. They were transferred to a −20C freezer 24 hours prior to tissue collection. Using chilled single-edged blades, the cerebellum was removed prior to placing the brain cortex-side up in a coronal rat brain matrix (Harvard Apparatus, Holliston, MA). Two mm-thick cortical tissue sections (approximately 4.5-2.5 mm anterior to bregma) were cut with razor blades and 2.0 mm-diameter medial prefrontal cortical tissue punches were collected on dry ice using Integra Miltex Biopsy Punchers with Plunger System (Fisher Scientific, Waltham, MA). The punches were placed in labeled, pre-chilled 2 ml Pre-Filled Bead Tubes with 2.8 mm Ceramic Beads (VWR, Radnor, PA), and weighed. Samples were homogenized in 250 µL of 1x PBS using a Bead Mill Homogenizer (VWR, Radnor, PA) at a speed of 4.85 m/s set for two 20s cycles at an interval of 11s. An additional 250 µL of Lysis Buffer 17 (R&D Systems, Minneapolis, MN) and 2.5 µL of 100X Halt™ Protease and Phosphatase Inhibitor Cocktail (Thermo Fisher Scientific, Waltham, MA) were added, followed by 30 min of gentle agitation and 10 min of centrifugation (500g, 40C) before the supernatant was collected. Total BDNF Standard was reconstituted in 1ml milliQ water and Calibrator Diluent RD5K (450μL). This stock solution was used to produce a dilution series with the following concentrations (pg/mL): 1000, 500, 250, 125, 62.5, 31.3, 15.6, 0. BDNF standards and tissue protein levels were run in triplicate per manufacturer instructions using Total BDNF Quantikine ELISA Kit (Catalog #DBNT00, R&D Systems, Minneapolis, MN), a sandwich enzyme immunoassay technique. Optical density of each sample was read at 450 nm with a wavelength correction set to 540 nm using a BioTek Gen5 Microplate Reader and Imager Software (Agilent, Santa Clara, CA). The triplicate OD measurements for each standard and tissue sample were averaged and the BDNF concentration in pg/ml was calculated and converted to pg/mg tissue weight.

### Experimental Design and Statistical Analyses

All statistical analyses were performed in GraphPad Prism 7.0.5 (GraphPad, USA). A Shapiro–Wilk test was used to assess the normal distribution of data.

Experiment 1: We compared active- and inactive lever presses, respectively, during both the 10-day self-administration period as well as the 10-day extinction period in sham-stimulated (n = 10) and VNS-treated rats (n = 11) using separate two-way ANOVAs with the factors group and time. A one-way ANOVA was performed on the cue-induced reinstatement session. Post-hoc analyses of main effects and interactions used Tukey’s multiple comparisons tests.

Experiment 2: We compared active-lever presses during the self-administration period using a two-way ANOVA with the factors time (sessions) and group (sham vehicle, n= 10; sham ANA12, n=11; VNS vehicle, n=13; VNS ANA-12, n= 9). Lever presses during the extinction period were analyzed using a separate three-way ANOVA with the factors stimulation (VNS vs. sham), drug (vehicle vs. ANA-12), and time. A two-way ANOVA with the factors stimulation and drug was performed on the cue-induced reinstatement session. Post-hoc analyses of main effects and interactions were performed using Tukey’s multiple comparisons tests.

Experiment 3: Single session extinction data and BDNF ELISA data for sham-stimulated (n = 11) VNS-treated rats (n = 12) were compared using unpaired two-tailed t-tests.

Experiment 4: We compared AMPAR:NMDAR ratios in 3 groups (naïve, 8 Recordings from 4 rats; yoked-saline, 9 recordings from 7 rats; cocaine, 10 recordings from 8 rats) using a one-way ANOVA. Post-hoc analysis used Tukey’s multiple comparisons test.

Experiment 5: Data obtained in this experiment stemmed from a subset of the rats used in experiment 2. We compared AMPAR:NMDAR ratios (as well as the changes in the individual currents) in 4 groups (sham vehicle, 8 cells from 4 rats; sham ANA12, 6 cells from 4 rats; VNS vehicle, 8 cells from 6 rats; VNS ANA-12, 8 cells from 7 rats) using separate two-way ANOVAs with the factors stimulation (sham or VNS) and drug (vehicle or ANA-12). Post-hoc analyses of main effects and interactions were performed using Tukey’s multiple comparisons tests. The correlation between behavior and AMPAR:NMDAR ratios was compared with a simple linear regression.

An α level of p<0.05 was considered significant in all experiments. Data are expressed as mean ± standard error of the mean (SEM). Individual responses are plotted over averaged responses. Asterisks on all figures represent differences between groups as indicated in the figure legends. Experimental designs and sample sizes are aimed at minimizing usage of animals and are sufficient for detecting robust effect sizes. While no statistical methods were used to predetermine sample sizes, our sample sizes are similar to those reported in previous publications.

## Results

### Experiment 1: Vagus nerve stimulation facilitates extinction learning and reduces cue-induced reinstatement

In order to determine the effects of VNS on drug-seeking behavior, rats were trained to self-administer cocaine for 15-18 days, followed by 10 days of extinction paired with VNS (*n* = 11) or sham-stimulation (*n* = 10). After 10 days, reinstatement to drug-seeking was measured in a cued reinstatement session by presenting the conditioned drug cues (**Figure 1**). A two-way ANOVA with the factors group and time (session) showed that that there were no differences in the rates of responding on the active lever between groups over the last 10 days of drug self-administration (group, F_(1,19)_ = 0.021, p = 0.886; time, F_(9,171)_ = 0.49, p = 0.88), and responses on the active lever exceeded responses on the inactive lever in both groups (two-way ANOVAs; sham, F_(1,18)_ = 37.92, p < 0.0001; VNS, F_(1,20)_ = 37.05, p < 0.0001). We used responses at the previously active lever as an indicator of extinction learning. A repeated-measures two-way ANOVA with the factors time (sessions) and treatment (Sham or VNS) showed a significant degree of extinction (main effect of time, F_(9,170)_ = 12.91, p < 0.0001), a significant difference between treatment groups (main effect of treatment, F_(1,19)_ = 17.48, p = 0.0005, **Figure 1A**), and an interaction between the factors (F_(9,171)_ = 3.26, p = 0.0011). Post-hoc analysis using Tukey’s multiple comparisons test showed that sham-simulated rats pressed the active lever significantly more on the first five days of extinction (day 1, p < 0.0001, **Figure 1B**; day 2, p = 0.0002; day 3, p = 0.0056; day 4, p = 0.0040; day 5, p = 0.0099). A separate two-way ANOVA for responses at the previously inactive lever showed significant treatment differences (F_(1,19)_ = 5.507, p = 0.03, **Figure 1A**). Post-hoc analysis with Tukey’s multiple comparisons test showed that sham-stimulated rats responded more often at the previously inactive lever on day 1 of extinction (p = 0.044, **Figure 1B**). Twenty-four hours after the last extinction session, drug-seeking was reinstated in a cued reinstatement session. Pairing extinction training with VNS significantly reduced responding at the active lever during cue-primed reinstatement (one-way ANOVA with active and inactive levers in sham- and VNS-treated rats; F_(3,38)_ = 13.16, p < 0.0001, **Figure 1C**). Post-hoc analysis with Tukey’s multiple comparisons test showed a significant decrease in active lever presses in VNS-treated rats (p = 0.0019), but no differences in responses at the inactive lever (p = 0.74).

**Figure 1:**
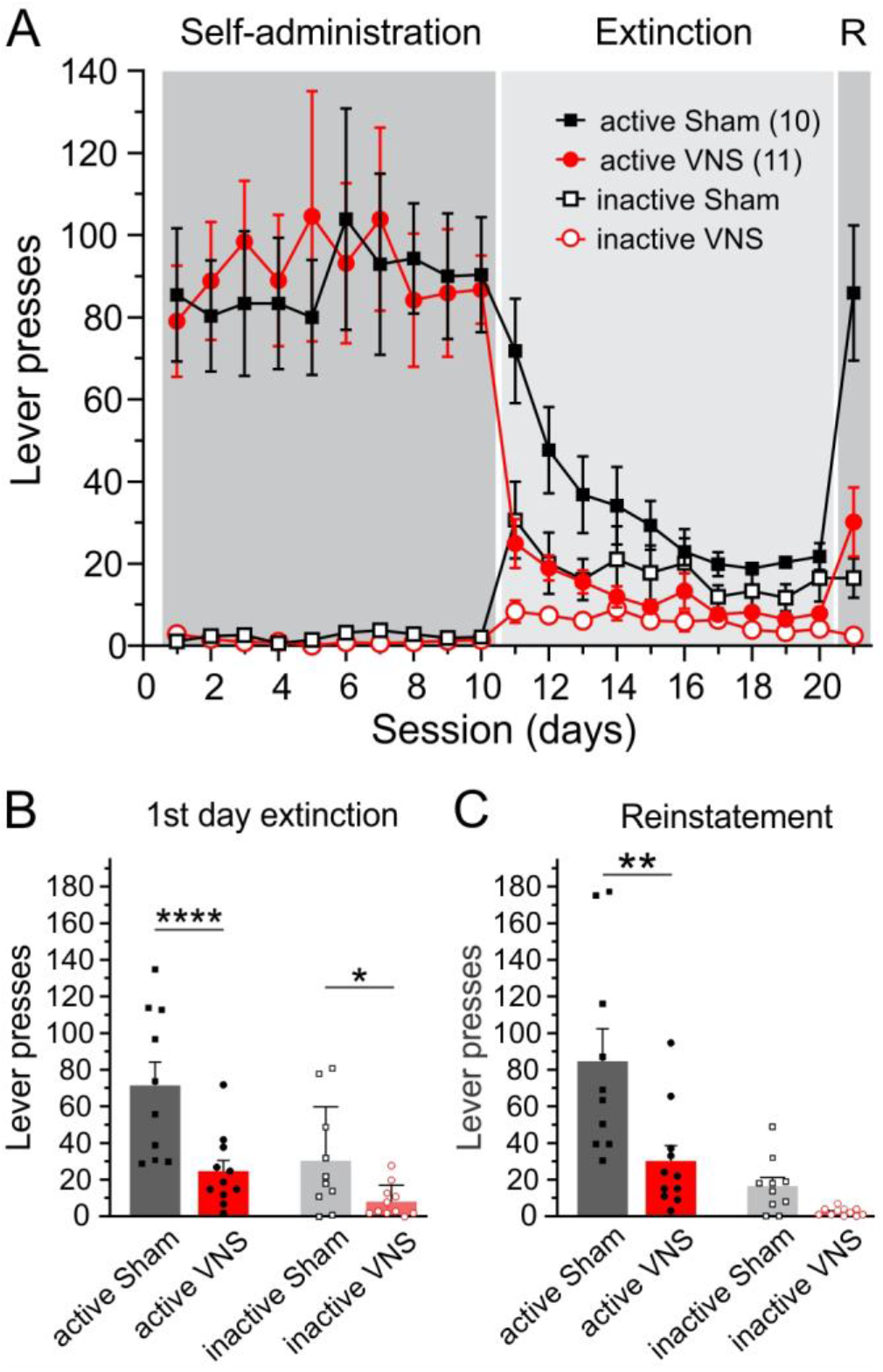
Vagus nerve stimulation (VNS) facilitates extinction from cocaine seeking and reduces cue-induced reinstatement. Rats were trained to self-administer cocaine and then underwent extinction training for 10 days while either receiving VNS (circles) or sham-stimulation (squares) before drug-seeking was reinstated by presentation of the previously drug-paired cues during a reinstatement session (R). A) Responses of VNS- or sham-stimulated rats at the active-(solid symbols) or inactive-(open symbols) lever during cocaine self-administration (last 10 days are shown), extinction, and cue-induced reinstatement. VNS was delivered during the 2-hr extinction sessions on days 11-20. B) Two-way ANOVAs over the extinction period show that rats which received VNS showed a reduction in both active and inactive lever presses during the first day of extinction. C) A one-ANOVA for the responses during the reinstatement session shows that VNS-treated rats also respond less at the active lever during cued-reinstatement (see text for details). P values are (*) <0.05, (**) <0.01, and (****) <0.0001.

### Experiment 2: Systemic blockade of TrkB receptors alters VNS’ effects on extinction and cue-induced reinstatement

VNS alters the levels of several neuromodulators in the CNS, including brain-derived neurotrophic factor (BDNF) (Dorr and Debonnel, 2006; Follesa et al., 2007; Manta et al., 2013; Nichols et al., 2011; Roosevelt et al., 2006). In order to determine if VNS-induced BDNF release modulates extinction learning and cue-induced reinstatement we gave VNS-treated or sham-stimulated rats systemic intraperitoneal injections of the Tropomyosin receptor kinase B (TrkB) receptor antagonist ANA-12 (0.5 mg/kg in DMSO, or DMSO-vehicle at a volume of 0.5 ml/kg) during the 10 days of extinction learning (but not during the reinstatement session) and examined their drug-seeking behavior (**Figure 2**). The combination of VNS or sham-stimulation and ANA-12 or vehicle infusion, respectively, resulted in 4 treatment groups (sham vehicle, n= 10; sham ANA12, n=11; VNS vehicle, n=13; VNS ANA-12, n= 9). A two-way ANOVA with the factors group and time showed no main effect of either factor and no interaction between the factors on active lever presses during the last ten days of self-administration (group, F_(3,39)_ = 0.195, p = 0.899; time, F_(9,351)_ = 0.775, p = 0.640; interaction, F_(27,351)_ = 0.805, p = 0.747), indicating that self-administration rates did not differ between groups prior to the start of extinction. For responses at the active lever during the extinction period a three-way ANOVA with the factors stimulation (Sham or VNS), drug (Vehicle or ANA-12) and time revealed main effects of stimulation (F_(1,39)_ = 6.91, p = 0.012) and time (F_(9,351)_ = 37.72, p<0.0001), but not drug (F_(1,39)_ = 0.057, p = 0.81), as well as interactions between stimulation and drug (F_(1,39)_ = 6.35, p = 0.016), and between all three factors (F_(9,351)_ = 6.15, p < 0.0001). Post hoc analysis showed that on the first day of extinction sham-vehicle treated rats responded significantly more on the previously active lever than VNS vehicle (p < 0.0001), sham ANA-12 (p < 0.0001), or VNS ANA-12 treated rats (p = 0.025) (**Figure 2B**). A separate two-way ANOVA with the factors stimulation and drug was performed on the active-lever responses during the cue-induced reinstatement session indicating a significant effect of stimulation (F_(1,39)_ = 4.99, p = 0.031), and an interaction of the factors (F_(1,39)_ = 13.55, p = 0.0007), but no effect of drug (F_(1,39)_ = 2.44, p = 0.126). Post hoc analysis with Tukey’s multiple comparisons test showed that VNS vehicle rats performed significantly fewer active lever presses than rats in all other groups (sham vehicle, p = 0.0006; sham ANA-12, p = 0.034; VNS ANA-12, p = 0.0034, **Figure 2C**). Taken together, these results show that TrkB receptor blockade alters the effects of VNS on extinction training and reinstatement, causing VNS ANA-12-treated rats to respond like sham-stimulated rats.

**Figure 2:**
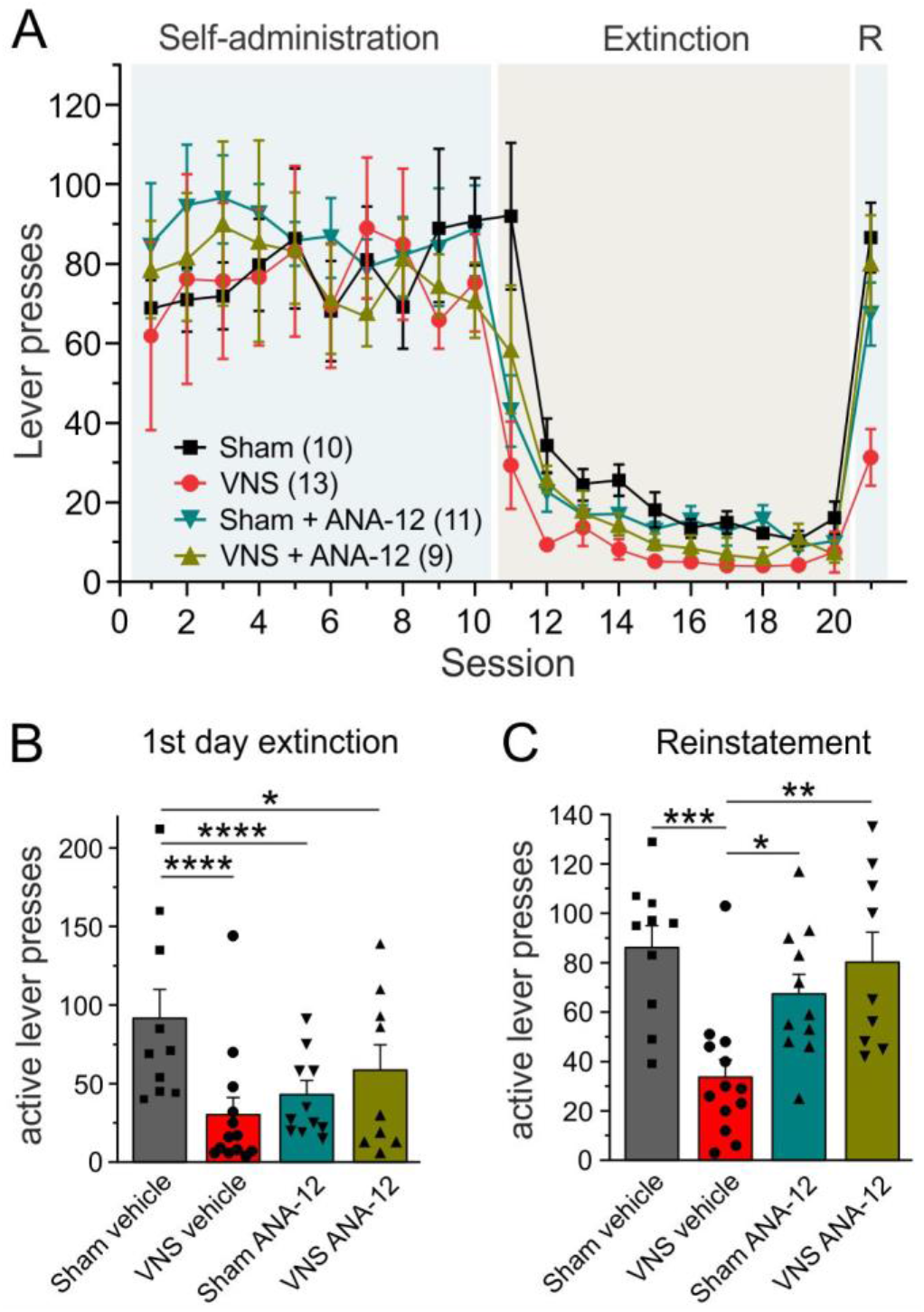
The effects of vagus nerve stimulation (VNS) on extinction learning and cue-induced reinstatement depend on activation of TrkB receptors. A) Active-lever presses of VNS- or sham-stimulated rats that received intraperitoneal infusions of either vehicle solution (sham, black squares; VNS, red circles) or the TrkB antagonist ANA-12 (sham, teal triangles; VNS, olive triangles) during extinction training on days 11-20. Shown are the last 10 days of cocaine-self-administration, extinction training, and cue-induced reinstatement (R). B) Results of a three-way ANOVA show that rats which received VNS and vehicle (red bar) showed the largest reduction in active lever presses during the first day of extinction. Rats in the sham ANA-12 and VNS ANA-12 groups also responded less than sham vehicle-treated rats. C) Results of a separate two-way ANOVA on the responses during the reinstatement session show that VNS-treated rats also show fewer active-lever presses during cue-induced reinstatement. Blocking TrkB receptors with ANA-12 abolished this effect, resulting in drug-seeking behavior similar to that seen in sham vehicle-treated rats. P values are (*) <0.05, (**) <0.01, (***) <0.001, and (****) <0.0001.

### Experiment 3: VNS during extinction training leads to elevated BDNF levels in the medial prefrontal cortex (mPFC)

Infusion of BDNF in the mPFC following cocaine self-administration reduces relapse to cocaine-seeking after abstinence, and reinstatement following extinction (Barry and McGinty, 2017; Berglind et al., 2007; Berglind et al., 2009; Go et al., 2016; Whitfield et al., 2011). VNS can increase BDNF levels in the brain (Follesa et al., 2007; Olsen et al., 2022), and thus we sought to determine if our conditions of VNS also lead to significant changes in BDNF levels that could explain its effects on extinction (**Figure 3**). Rats self-administered cocaine followed by one day of extinction training paired with sham-stimulation (n = 11) or VNS (n = 12) (**Figure 3A**). An unpaired t-test showed VNS-treated animals responded significantly less on the previously active lever during the extinction session (t_(21)_ = 3.583, p = 0.0018, **Figure 3B**). Immediately following the extinction session rats were killed for quantification of BDNF levels in mPFC via ELISA. Assays were performed in two separate cohorts and therefore measures are expressed as the percent change in BDNF levels compared to the average of the sham brains within each cohort. An unpaired t test showed VNS-treated rats had significantly higher levels of BDNF in the mPFC compared to sham rats (t_(21)_ = 2.452, p = 0.023, **Figure 3C**).

**Figure 3:**
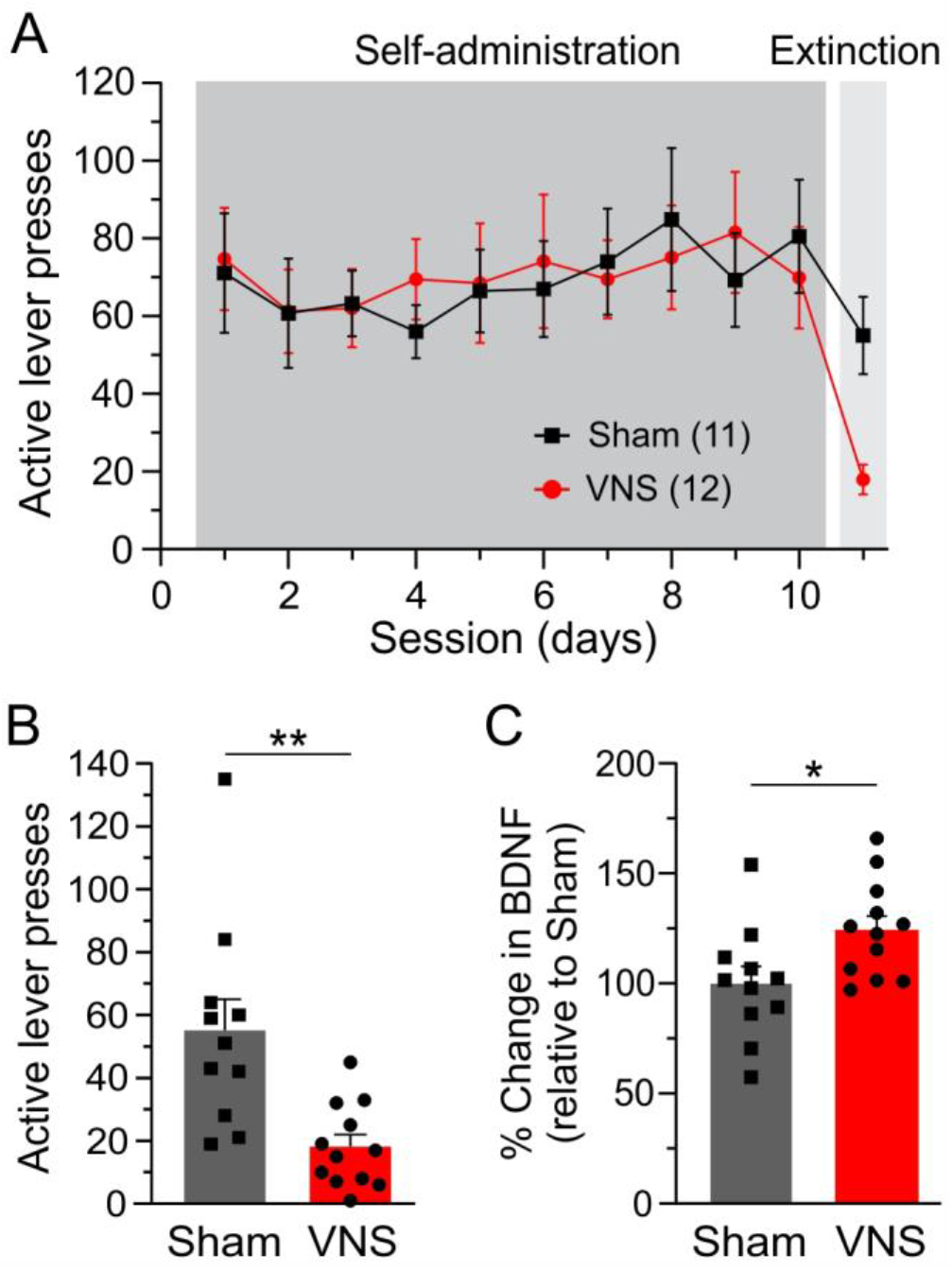
Vagus nerve stimulation (VNS) increases the levels of BDNF in the medial prefrontal cortex. A) Active-lever presses during the last 10 days of cocaine self-administration and a single day of extinction training during which rats received either sham-stimulation (black squares) or VNS (red circles). B) VNS reduced active lever presses during the extinction session. C) Pairing extinction training with VNS increased BDNF content in mPFC tissue punches as measured by ELISA. Results are expressed as percent change from the average of the sham-stimulated group across two different cohorts (see text for detail). Unpaired t-tests in B) and C). P values are (*) <0.05 and (**) <0.01.

### Experiment 4: Cocaine self-administration leads to alterations in the AMPAR:NMDAR current ratio in the mPFC

Cocaine alters glutamatergic transmission in the mPFC (Caffino et al., 2018; Kasanetz et al., 2013; Otis and Mueller, 2017; Sasase et al., 2019). Alterations in the AMPAR:NMDAR current ratio provide sensitive assays for changes in postsynaptic receptor function. We first examined the effects of cocaine self-administration on glutamate receptors in the IL. Figure **4A** shows active lever presses in a cohort of rats self-administering cocaine or receiving yoked saline infusions. **Figure 4B** shows representative voltage-clamp recordings of AMPA and NMDA currents from IL layer 5 pyramidal neurons from cocaine-administering or yoked-saline rats, killed immediately after their last self-administration session, as well as a group of drug naïve rats for comparison. A one-way ANOVA found a significant group effect in AMPAR:NMDAR ratios (F_(2,24)_ = 4.811, p = 0.017, **Figure 4C**), and post-hoc testing with Tukey’s multiple comparisons test indicated that AMPAR:NMDAR ratios in cocaine self-administering rats were significantly smaller than in both naïve rats (p = 0.03) and yoked-saline rats (p = 0.044). Ratios of AMPA and NMDA currents were similar in yoked-saline and naïve rats (p = 0.966).

**Figure 4:**
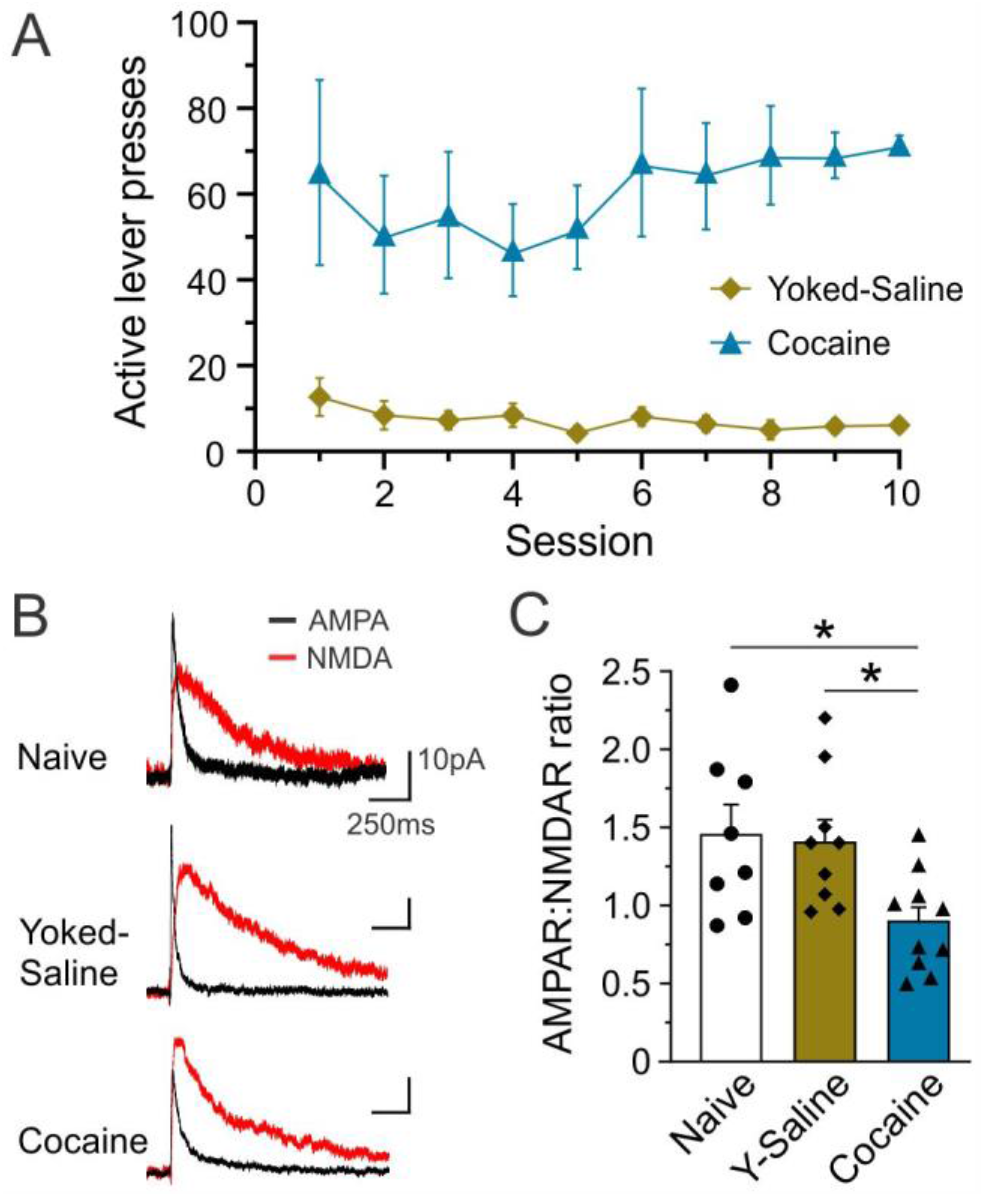
Cocaine self-administration alters the ratio of currents through AMPA- and NMDA receptors. A) Active-lever presses in rats self-administering cocaine (blue triangles) and their yoked-saline controls (diamonds) during the last 10 days of self-administration. Brain slices for whole-cell patch-clamp recordings were prepared immediately after the last self-administration session (day 10). B) Representative currents through AMPA-(black lines) and NMDA receptors (red lines) from layer 5 pyramidal cells in the infralimbic cortex (IL) of naïve rats, or rats that received yoked-saline or cocaine-infusions as illustrated in A). C) Results of a one-way ANOVA show that cocaine self-administration lowers the AMPAR:NMDAR ratio in IL pyramidal neurons compared to cells from naïve or yoked saline-treated rats. P values are (*) <0.05.

### Experiment 5: Pairing extinction with VNS reverses drug-induced changes in AMPAR:NMDAR ratios and is blocked by ANA-12

The effects of BDNF infusions into the mPFC on reinstatement are mediated by glutamatergic mechanisms (Barry and McGinty, 2017; Berglind et al., 2007; Berglind et al., 2009; Go et al., 2016; Whitfield et al., 2011). Therefore, we next determined whether pairing extinction with VNS a) can reverse cocaine-induced alterations in glutamatergic signaling, and b) whether TrkB receptor signaling is required for this effect. We obtained whole-cell recordings of AMPA and NMDA currents in layer 5 IL pyramidal neurons immediately following the reinstatement session in a subset of rats that received VNS or sham stimulation with ANA-12 or vehicle during extinction in experiment 2 (**Figure 5A**). A two-way ANOVA with the factors stimulation (Sham or VNS) and drug (vehicle or ANA-12) found significant main effects for both factors and an interaction of the effects on AMPAR:NMDAR ratios (stimulation, F_(1,26)_ = 4.689, p = 0.040; drug, F_(1,26)_ = 5.718, p = 0.024; interaction, F_(1,26)_ = 22.14, p < 0.0001). Post-hoc testing with Tukey’s multiple comparisons test showed that AMPAR:NMDAR ratios in recordings from VNS-vehicle rats were significantly larger than in recordings from sham-vehicle rats (p = 0.0001), sham-ANA-12 rats (p = 0.022), or VNS-ANA-12 rats (p = 0.0002). AMPAR:NMDAR ratios in sham-vehicle rats were not different from those in sham ANA-12 rats (p = 0.32), or rats in the VNS ANA-12 group (p > 0.998). Similarly, sham-ANA-12 rats and VNS-ANA-12 rats showed comparable AMPAR:NMDAR ratios (p = 0.41) (**Figure 5B**) These changes in synaptic plasticity were mostly driven by changes in AMPA currents: A two-way ANOVA comparing raw AMPA currents found a significant main effect of stimulation (F_(1,26)_ = 7.151, p < 0.0128), and a drug X stimulation interaction (F_(1,26)_ = 6.578, p = 0.016), but no main effect for drug (F_(1,26)_ = 2.433, p = 0.13). Post-hoc analysis with Tukey’s multiple comparisons test indicated that AMPA currents in VNS vehicle-treated rats were significantly larger than in sham-vehicle rats (p = 0.0036), sham ANA-12 rats (p = 0.037), and VNS ANA-12 rats (p = 0.026) (**Figure 5C**). A similar comparison for NMDA currents in the same cells found no difference for either factor (two-way ANOVA; stimulation, F_(1,26)_ = 1.272, p = 0.2693; drug, F_(1,26)_ = 0.0000349, p = 0.995; interaction, F_(1,26)_ = 0.131, p = 0.720; **Figure 5D**). In order to determine whether VNS-induced BDNF modulation of glutamatergic synaptic transmission is correlated with relapse behavior we performed a simple linear regression. For each rat we averaged the AMPAR:NMDAR ratios across multiple recordings to obtain a single value for each animal and then correlated this ratio with the animal’s active lever presses during reinstatement. The regression analysis showed that AMPAR:NMDAR ratios in IL pyramidal neurons were highly correlated with reinstatement behavior (F_(1,19)_ = 16.92, p = 0.0006, r^2^ = 0.4711, **Figure 5E**), with higher AMPAR:NMDAR ratios associated with fewer lever presses. Taken together, our results suggest that pairing extinction with VNS reverses changes in AMPAR:NMDAR ratios in IL layer 5 pyramidal neurons caused by cocaine self-administration, and that this reversal is abolished by TrkB receptor blockade. Additionally, changes in glutamatergic signaling in the IL are correlated with drug-seeking during cue-induced reinstatement.

**Figure 5:**
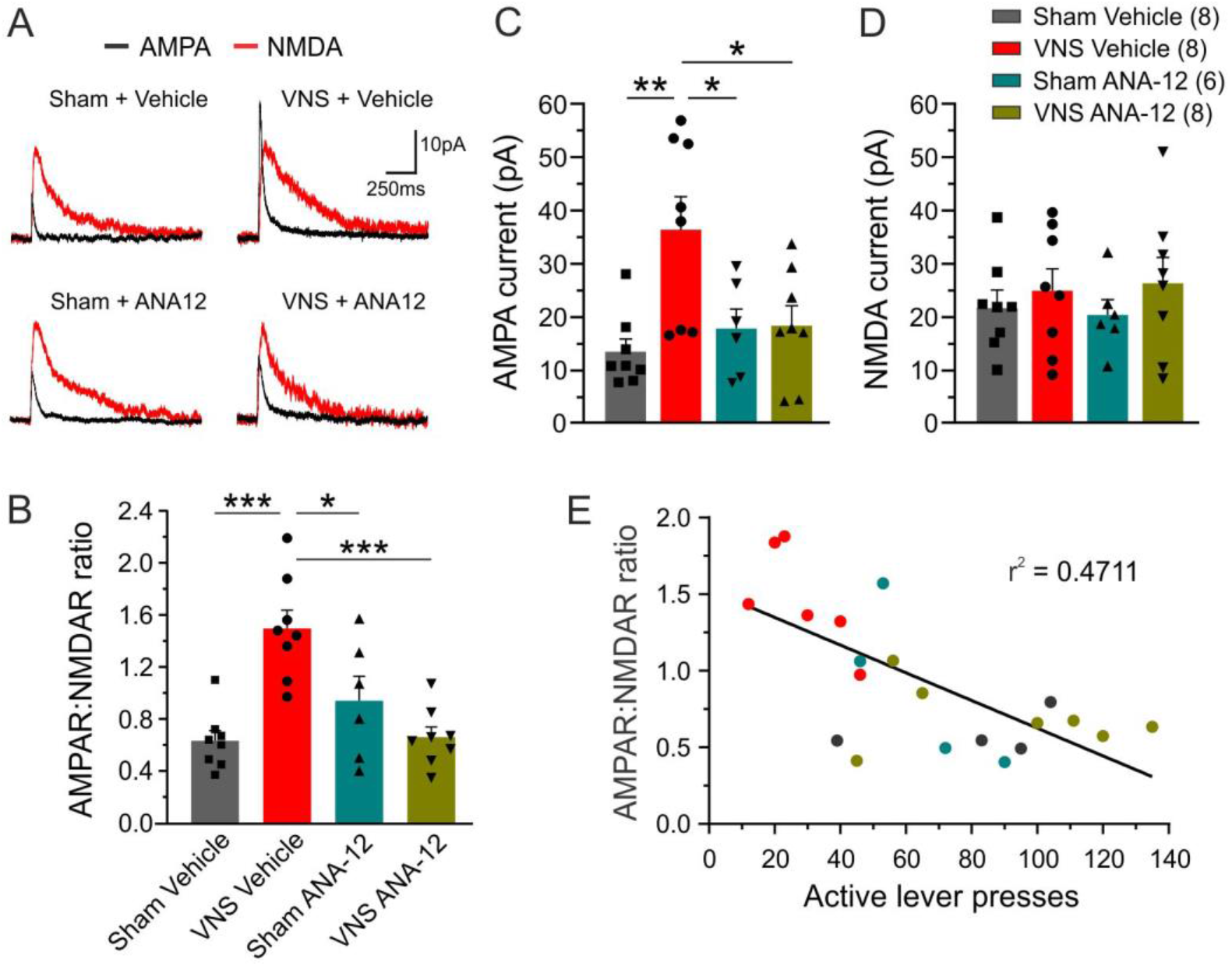
Vagus nerve stimulation (VNS) reverses cocaine-induced changes in glutamatergic plasticity, which is dependent on TrkB receptor activation. A) Representative currents through AMPA-(black lines) and NMDA receptors (red lines) from layer 5 pyramidal cells in the infralimbic cortex (IL) of rats that received intraperitoneal infusions of either vehicle solution or the TrkB antagonist ANA-12 during extinction training paired with VNS (VNS vehicle, VNS ANA-12) or sham stimulation (sham vehicle, sham ANA-12). B) A two-way ANOVA showed that IL pyramidal neurons from rats in the VNS vehicle group (red bar) have significant larger AMPAR:NMDAR ratios following cue-induced reinstatement than cells from sham vehicle-(grey bar), sham ANA-12-(teal bar), or VNS ANA-12-treated rats (olive bar). C, D) Two-way ANOVAs indicate that the shift in AMPAR:NMDAR ratios is driven by changes in AMPA currents, C), while the amplitudes of NMDAR currents did not significantly differ between the 4 treatment groups, D). E) A linear regression analysis correlating AMPAR:NMDAR ratios with drug-seeking behavior during reinstatement indicates that increased lever presses are correlated with lower AMPAR:NMDAR ratios. For some rats multiple recordings were obtained; in these cases, the ratios were averaged to correlate a single value with the number of active lever presses. P values are (*) <0.05 and (**) <0.01, and (***) <0.001.

## Discussion

Vagus nerve stimulation provides a means to modulate extinction circuits to improve treatment retention and outcome. Here we replicate previous findings which show that VNS facilitates extinction learning and reduces cue-induced reinstatement of cocaine-seeking (Childs et al., 2017; Childs et al., 2019), and we determine the importance of BDNF in these effects. VNS increased BDNF levels in the mPFC and systemic blockade of TrkB receptors diminished the effects of VNS on reinstatement. Cocaine self-administration altered glutamatergic synaptic plasticity in the IL, and pairing extinction with VNS reversed these effects in a TrkB-dependent manner that correlated with reinstatement behavior.

### VNS facilitates extinction learning

Stimulation of ascending fibers of the vagus nerve leads to the release of several neuromodulators, including norepinephrine, acetylcholine, and BDNF in the CNS that cause widespread cortical and subcortical activation (Cao et al., 2016; Collins et al., 2021). The release of neuromodulators alters cognitive and motivational states, in turn changing how contexts are perceived and remembered. Thus, VNS-induced changes may act as a primer, modulating synaptic plasticity in response to specific inputs that occur during sensory stimulation (Borland et al., 2018), or learning and memory (Pena et al., 2014; Porter et al., 2012). First, we replicated previous findings (Childs et al., 2017; Childs et al., 2019) to show that pairing extinction training with VNS 1.) accelerates extinction learning during the initial session, and 2.) decreases cue-induced reinstatement, suggesting enhanced consolidation of extinction memories. Previous work has shown that VNS does not affect behavior during the extinction training period by interrupting ongoing behavior or acting as punishment (Childs et al., 2017). Stimulation of the left vagus nerve also does not induce conditioned place preference or avoidance, indicating that by itself it is neither rewarding or aversive (Childs et al., 2019; Noble et al., 2019a).

### The importance of BDNF for the effects of VNS on extinction learning

Dysregulation of BDNF is associated with the establishment and maintenance of psychostimulant-induced memories, as well as the mechanisms that underlie extinction learning. Chronic cocaine exposure causes a decrease in BDNF in several brain regions that may cause impaired neuroplasticity and reduced stress resilience, two factors that contribute to addiction (McGinty et al., 2010; Pitts et al., 2016; Verheij et al., 2016). The vagus nerve regulates constitutive BDNF expression in the brain (Amagase et al., 2023) and VNS has been shown to increase BDNF in various brain areas (Follesa et al., 2007; Olsen et al., 2022). We used the TrkB receptor antagonist ANA-12 to determine the contribution of BDNF to the effects of VNS on extinction and reinstatement. Systemic application of ANA-12 diminished the ability of VNS to facilitate extinction learning on the first day and to reduce cue-induced reinstatement. However, surprisingly, rats in the sham ANA-12 group also showed significantly reduced responding at the previously active lever during the first day of extinction, similar to VNS-treated rats. This effect might be explained by the rapid anxiolytic-like effects that ANA-12 has in mice (Cazorla et al., 2011), which might counter cocaine withdrawal-induced anxiety or uncertainty induced by the change in contingencies. However, rats in the sham ANA-12 group showed no reduction in responding during reinstatement, which suggests that the initial facilitation of extinction is not sufficient to reduce reinstatement 11 days later. The differential response rates during reinstatement also imply that the effect of VNS during (prolonged) extinction are likely not solely due to potential anxiolytic effects of VNS (Mathew et al., 2020; Noble et al., 2019b), but reflect changes in extinction learning. Aside from the changes on the first day, systemic ANA-12 did not significantly alter the trajectory or overall effectiveness of extinction training in sham-stimulated rats, suggesting that TrkB stimulation is not a necessary requirement for normal extinction learning.

In order to understand the mechanisms through which VNS and BDNF modulate extinction and reinstatement we focused on the mPFC, and specifically the IL, because of its importance in the acquisition and expression of extinction memories for drug-seeking (Augur et al., 2016; Peters et al., 2009). Repeated exposure to cocaine and withdrawal lower the levels of BDNF in the mPFC (Fumagalli et al., 2007; Fumagalli et al., 2013; McGinty et al., 2010). Extinction by itself leads to a small increase in BDNF in the PFC (Hastings et al., 2020; Li et al., 2018). Our ELISA data from mPFC tissue obtained following a single extinction session show that VNS increases BDNF levels significantly above levels induced by extinction alone and that this increase correlates with facilitated extinction behavior. Taken together, our data suggest that pairing VNS with extinction increases the release of BDNF in the mPFC and that this release is crucial for the modulation of both extinction and reinstatement behavior. These results mimic previous findings which have shown that a single infusion of BDNF into the prelimbic cortex (PL) directly after a final cocaine self-administration session reduces relapse to cocaine-seeking after abstinence, as well as cue- and cocaine prime-induced reinstatement following extinction (Barry and McGinty, 2017; Berglind et al., 2007; Berglind et al., 2009; Go et al., 2016; Whitfield et al., 2011), while infralimbic BDNF can facilitate extinction of cocaine-seeking (Otis et al., 2014; Yousuf et al., 2019). The effects of VNS are difficult to compare to those of a single high dose of BDNF infused into the mPFC because of differences in dosage and time course that affect BDNF’s action. Evidence from sensory and motor learning suggests that the temporally precise modulation of active networks (in our case networks actively engaged in extinction learning) greatly contributes to the effectiveness of VNS (Borland et al., 2018; Hays et al., 2014; Ruiz et al., 2023). Thus, this network-specificity may compensate for the relatively smaller physiological increases in BDNF produced by VNS. However, our current data does not address the question of how often VNS-induced BDNF release has to be paired with extinction to effectively reduce reinstatement.

### Cocaine and VNS-induced alterations in synaptic plasticity

Cocaine induces aberrant plasticity at glutamatergic synapses in the mPFC, leading to altered salience attribution and cue-induced drug-seeking (Caffino et al., 2018; Kasanetz et al., 2013; Otis and Mueller, 2017). The direction and magnitude of changes depends on the brain region examined, and the timepoint (acute vs. prolonged withdrawal, extinction, or after reinstatement) at which effects are tested. Cocaine self-administration leads to a decrease in NMDA receptor expression, as well as GluN2B dephosphorylation in the mPFC during withdrawal (Ben-Shahar et al., 2007; Sun et al., 2013). However, previous work in the PL has also shown increases in AMPA receptor function following cocaine regimen that lead to locomotor sensitization (Ruan and Yao, 2021) or conditioned place preference (Otis and Mueller, 2017). Here we show that AMPAR:NMDAR ratios in IL pyramidal neurons are reduced during acute cocaine withdrawal. Similarly, sham-stimulated rats showed reduced AMPAR:NMDAR ratios after relapse, which were inversely correlated with the rate of responding at the active lever during the cue-induced reinstatement session. These shifts in synaptic plasticity were due to smaller AMPA currents in sham-stimulated rats. Taken together, these findings suggest that cocaine self-administration and reinstatement alter synaptic plasticity in the IL by reducing AMPA currents. In contrast, in rats that received VNS during extinction training the AMPAR:NMDAR current ratios following cued reinstatement closely resembled those of naïve or YS rats. Therefore, VNS modulates glutamatergic plasticity in the IL to reduce reinstatement, presumably by strengthening effects that occur during extinction learning. Importantly, our results show that these effects depend on stimulation of TrkB receptors. As detailed above, infusions of BDNF into the PL reduce relapse to cocaine-seeking after abstinence, as well as cue-induced reinstatement following extinction. Inhibiting TrkB receptors, ERK/MAP-kinase activation, or NMDA receptors blocks these effects (Barry and McGinty, 2017; Berglind et al., 2007; Berglind et al., 2009; Go et al., 2016; Whitfield et al., 2011), indicating an important interaction between TrkB signaling and NMDARs in BDNF’s suppressive effects on drug-seeking (Otis et al., 2014; Otis and Mueller, 2017). However, BDNF can also mediate activity-dependent changes in synaptic strength by regulating AMPAR trafficking (Fukumoto et al., 2020; Li and Wolf, 2011, 2015; Reimers et al., 2014). In our experiments, TrkB receptor blockade during extinction training primarily affected the VNS-modulation of AMPA currents, resulting in reduced AMPAR:NMDAR ratios in the ANA-12-treated groups during reinstatement that paralleled those in the sham-treated group.

Extinguishing reactivity to drug-associated environments and extinguishing the instrumental response for drugs are both important for the treatment of SUD (Childress et al., 1993). Progress made during rehab is often undone when patients return to the environments formerly associated with drug-taking (Hammond and Wagner, 2013). Our data add further support for the idea that VNS can be used to promote synaptic plasticity to increase extinction from drug-seeking and to reduce relapse, thus increasing the efficacy of behavioral therapies where pharmacological approaches have yielded mixed results (Kantak and Nic Dhonnchadha, 2011; Nic Dhonnchadha and Kantak, 2011). We also provide evidence that at least in the mPFC these changes are mediated by BDNF activation of TrkB receptors, which may further aid the development of adjunct treatments.

## COI statement

The authors declare no competing financial interests.

## Acknowledgements

This work was supported by grant 1R01 DA055008-01 to SK.

## Notes

### Competing Interest Statement

The authors have declared no competing interest.

## References

Amagase, Y., Kambayashi, R., Sugiyama, A., Takei, Y., 2023. Peripheral Regulation of Central Brain-Derived Neurotrophic Factor Expression through the Vagus Nerve. Int J Mol Sci 24.

Augur, I.F., Wyckoff, A.R., Aston-Jones, G., Kalivas, P.W., Peters, J., 2016. Chemogenetic Activation of an Extinction Neural Circuit Reduces Cue-Induced Reinstatement of Cocaine Seeking. J Neurosci 36, 10174–10180.

Barry, S.M., McGinty, J.F., 2017. Role of Src Family Kinases in BDNF-Mediated Suppression of Cocaine-Seeking and Prevention of Cocaine-Induced ERK, GluN2A, and GluN2B Dephosphorylation in the Prelimbic Cortex. Neuropsychopharmacology 42, 1972–1980.

Ben-Menachem, E., Manon-Espaillat, R., Ristanovic, R., Wilder, B.J., Stefan, H., Mirza, W., Tarver, W.B., Wernicke, J.F., 1994. Vagus nerve stimulation for treatment of partial seizures: 1. A controlled study of effect on seizures. First International Vagus Nerve Stimulation Study Group. Epilepsia 35, 616–626.

Ben-Shahar, O., Keeley, P., Cook, M., Brake, W., Joyce, M., Nyffeler, M., Heston, R., Ettenberg, A., 2007. Changes in levels of D1, D2, or NMDA receptors during withdrawal from brief or extended daily access to IV cocaine. Brain Res 1131, 220–228.

Berglind, W.J., See, R.E., Fuchs, R.A., Ghee, S.M., Whitfield, T.W., Jr., Miller, S.W., McGinty, J.F., 2007. A BDNF infusion into the medial prefrontal cortex suppresses cocaine seeking in rats. Eur J Neurosci 26, 757–766.

Berglind, W.J., Whitfield, T.W., Jr., LaLumiere, R.T., Kalivas, P.W., McGinty, J.F., 2009. A single intra-PFC infusion of BDNF prevents cocaine-induced alterations in extracellular glutamate within the nucleus accumbens. J Neurosci 29, 3715–3719.

Borland, M.S., Engineer, C.T., Vrana, W.A., Moreno, N.A., Engineer, N.D., Vanneste, S., Sharma, P., Pantalia, M.C., Lane, M.C., Rennaker, R.L., Kilgard, M.P., 2018. The Interval Between VNS-Tone Pairings Determines the Extent of Cortical Map Plasticity. Neuroscience 369, 76–86.

Caffino, L., Messa, G., Fumagalli, F., 2018. A single cocaine administration alters dendritic spine morphology and impairs glutamate receptor synaptic retention in the medial prefrontal cortex of adolescent rats. Neuropharmacology 140, 209–216.

Cao, B., Wang, J., Shahed, M., Jelfs, B., Chan, R.H., Li, Y., 2016. Vagus Nerve Stimulation Alters Phase Synchrony of the Anterior Cingulate Cortex and Facilitates Decision Making in Rats. Sci Rep 6, 35135.

Cazorla, M., Premont, J., Mann, A., Girard, N., Kellendonk, C., Rognan, D., 2011. Identification of a low-molecular weight TrkB antagonist with anxiolytic and antidepressant activity in mice. J Clin Invest 121, 1846–1857.

Childress, A.R., Hole, A.V., Ehrman, R.N., Robbins, S.J., McLellan, A.T., O’Brien, C.P., 1993. Cue reactivity and cue reactivity interventions in drug dependence. NIDA Res Monogr 137, 73–95.

Childs, J.E., Alvarez-Dieppa, A.C., McIntyre, C.K., Kroener, S., 2015. Vagus Nerve Stimulation as a Tool to Induce Plasticity in Pathways Relevant for Extinction Learning. J Vis Exp, e53032.

Childs, J.E., DeLeon, J., Nickel, E., Kroener, S., 2017. Vagus nerve stimulation reduces cocaine seeking and alters plasticity in the extinction network. Learn Mem 24, 35–42.

Childs, J.E., Kim, S., Driskill, C.M., Hsiu, E., Kroener, S., 2019. Vagus nerve stimulation during extinction learning reduces conditioned place preference and context-induced reinstatement of cocaine seeking. Brain Stimul 12, 1448–1455.

Clark, K.B., Krahl, S.E., Smith, D.C., Jensen, R.A., 1995. Post-training unilateral vagal stimulation enhances retention performance in the rat. Neurobiol Learn Mem 63, 213–216.

Clark, K.B., Naritoku, D.K., Smith, D.C., Browning, R.A., Jensen, R.A., 1999. Enhanced recognition memory following vagus nerve stimulation in human subjects. Nat Neurosci 2, 94–98.

Collins, L., Boddington, L., Steffan, P.J., McCormick, D., 2021. Vagus nerve stimulation induces widespread cortical and behavioral activation. Curr Biol 31, 2088–2098 e2083.

Conklin, C.A., Tiffany, S.T., 2002. Applying extinction research and theory to cue-exposure addiction treatments. Addiction 97, 155–167.

Dawson, J., Liu, C.Y., Francisco, G.E., Cramer, S.C., Wolf, S.L., Dixit, A., Alexander, J., Ali, R., Brown, B.L., Feng, W., DeMark, L., Hochberg, L.R., Kautz, S.A., Majid, A., O’Dell, M.W., Pierce, D., Prudente, C.N., Redgrave, J., Turner, D.L., Engineer, N.D., Kimberley, T.J., 2021. Vagus nerve stimulation paired with rehabilitation for upper limb motor function after ischaemic stroke (VNS-REHAB): a randomised, blinded, pivotal, device trial. Lancet 397, 1545–1553.

Dorr, A.E., Debonnel, G., 2006. Effect of vagus nerve stimulation on serotonergic and noradrenergic transmission. J Pharmacol Exp Ther 318, 890–898.

Ehrman, R.N., Robbins, S.J., Childress, A.R., O’Brien, C.P., 1992. Conditioned responses to cocaine-related stimuli in cocaine abuse patients. Psychopharmacology (Berl) 107, 523–529.

Follesa, P., Biggio, F., Gorini, G., Caria, S., Talani, G., Dazzi, L., Puligheddu, M., Marrosu, F., Biggio, G., 2007. Vagus nerve stimulation increases norepinephrine concentration and the gene expression of BDNF and bFGF in the rat brain. Brain Res 1179, 28–34.

Fowler, J.S., Volkow, N.D., Kassed, C.A., Chang, L., 2007. Imaging the addicted human brain. Sci Pract Perspect 3, 4–16.

Fukumoto, K., Fogaca, M.V., Liu, R.J., Duman, C.H., Li, X.Y., Chaki, S., Duman, R.S., 2020. Medial PFC AMPA receptor and BDNF signaling are required for the rapid and sustained antidepressant-like effects of 5-HT(1A) receptor stimulation. Neuropsychopharmacology 45, 1725–1734.

Fumagalli, F., Di Pasquale, L., Caffino, L., Racagni, G., Riva, M.A., 2007. Repeated exposure to cocaine differently modulates BDNF mRNA and protein levels in rat striatum and prefrontal cortex. Eur J Neurosci 26, 2756–2763.

Fumagalli, F., Moro, F., Caffino, L., Orru, A., Cassina, C., Giannotti, G., Di Clemente, A., Racagni, G., Riva, M.A., Cervo, L., 2013. Region-specific effects on BDNF expression after contingent or non-contingent cocaine i.v. self-administration in rats. Int J Neuropsychopharmacol 16, 913–918.

Go, B.S., Barry, S.M., McGinty, J.F., 2016. Glutamatergic neurotransmission in the prefrontal cortex mediates the suppressive effect of intra-prelimbic cortical infusion of BDNF on cocaine-seeking. Eur Neuropsychopharmacol 26, 1989–1999.

Gutman, A.L., Nett, K.E., Cosme, C.V., Worth, W.R., Gupta, S.C., Wemmie, J.A., LaLumiere, R.T., 2017. Extinction of Cocaine Seeking Requires a Window of Infralimbic Pyramidal Neuron Activity after Unreinforced Lever Presses. J Neurosci 37, 6075–6086.

Hammond, S., Wagner, J.J., 2013. The context dependency of extinction negates the effectiveness of cognitive enhancement to reduce cocaine-primed reinstatement. Behav Brain Res 252, 444–449.

Hastings, M.H., Gauthier, J.M., Mabry, K., Tran, A., Man, H.Y., Kantak, K.M., 2020. Facilitative effects of environmental enrichment for cocaine relapse prevention are dependent on extinction training context and involve increased TrkB signaling in dorsal hippocampus and ventromedial prefrontal cortex. Behav Brain Res 386, 112596.

Hays, S.A., Khodaparast, N., Ruiz, A., Sloan, A.M., Hulsey, D.R., Rennaker, R.L., 2nd, Kilgard, M.P., 2014. The timing and amount of vagus nerve stimulation during rehabilitative training affect poststroke recovery of forelimb strength. Neuroreport 25, 676–682.

Kantak, K.M., Nic Dhonnchadha, B.A., 2011. Pharmacological enhancement of drug cue extinction learning: translational challenges. Ann N Y Acad Sci 1216, 122–137.

Kasanetz, F., Lafourcade, M., Deroche-Gamonet, V., Revest, J.M., Berson, N., Balado, E., Fiancette, J.F., Renault, P., Piazza, P.V., Manzoni, O.J., 2013. Prefrontal synaptic markers of cocaine addiction-like behavior in rats. Mol Psychiatry 18, 729–737.

Li, C., White, A.C., Schochet, T., McGinty, J.F., Frantz, K.J., 2018. ARC and BDNF expression after cocaine self-administration or cue-induced reinstatement of cocaine seeking in adolescent and adult male rats. Addict Biol 23, 1233–1241.

Li, X., Wolf, M.E., 2011. Brain-derived neurotrophic factor rapidly increases AMPA receptor surface expression in rat nucleus accumbens. Eur J Neurosci 34, 190–198.

Li, X., Wolf, M.E., 2015. Multiple faces of BDNF in cocaine addiction. Behav Brain Res 279, 240–254.

Liu, H., Liu, Y., Yu, J., Lai, M., Zhu, H., Sun, A., Chen, W., Zhou, W., 2011. Vagus nerve stimulation inhibits heroin-seeking behavior induced by heroin priming or heroin-associated cues in rats. Neurosci Lett 494, 70–74.

Liu, J., Liang, J., Qin, W., Tian, J., Yuan, K., Bai, L., Zhang, Y., Wang, W., Wang, Y., Li, Q., Zhao, L., Lu, L., von Deneen, K.M., Liu, Y., Gold, M.S., 2009. Dysfunctional connectivity patterns in chronic heroin users: an fMRI study. Neurosci Lett 460, 72–77.

Manta, S., El Mansari, M., Debonnel, G., Blier, P., 2013. Electrophysiological and neurochemical effects of long-term vagus nerve stimulation on the rat monoaminergic systems. Int J Neuropsychopharmacol 16, 459–470.

Mathew, E., Tabet, M.N., Robertson, N.M., Hays, S.A., Rennaker, R.L., Kilgard, M.P., McIntyre, C.K., Souza, R.R., 2020. Vagus nerve stimulation produces immediate dose-dependent anxiolytic effect in rats. J Affect Disord 265, 552–557.

McGinty, J.F., Whitfield, T.W., Jr., Berglind, W.J., 2010. Brain-derived neurotrophic factor and cocaine addiction. Brain Res 1314, 183–193.

Millan, E.Z., Marchant, N.J., McNally, G.P., 2011. Extinction of drug seeking. Behav Brain Res 217, 454–462.

Muller Ewald, V.A., De Corte, B.J., Gupta, S.C., Lillis, K.V., Narayanan, N.S., Wemmie, J.A., LaLumiere, R.T., 2019. Attenuation of cocaine seeking in rats via enhancement of infralimbic cortical activity using stable step-function opsins. Psychopharmacology (Berl) 236, 479–490.

Nic Dhonnchadha, B.A., Kantak, K.M., 2011. Cognitive enhancers for facilitating drug cue extinction: insights from animal models. Pharmacology, biochemistry, and behavior 99, 229–244.

Nichols, J.A., Nichols, A.R., Smirnakis, S.M., Engineer, N.D., Kilgard, M.P., Atzori, M., 2011. Vagus nerve stimulation modulates cortical synchrony and excitability through the activation of muscarinic receptors. Neuroscience 189, 207–214.

Noble, L.J., Chuah, A., Callahan, K.K., Souza, R.R., McIntyre, C.K., 2019a. Peripheral effects of vagus nerve stimulation on anxiety and extinction of conditioned fear in rats. Learn Mem 26, 245–251.

Noble, L.J., Meruva, V.B., Hays, S.A., Rennaker, R.L., Kilgard, M.P., McIntyre, C.K., 2019b. Vagus nerve stimulation promotes generalization of conditioned fear extinction and reduces anxiety in rats. Brain Stimul 12, 9–18.

Olsen, L.K., Moore, R.J., Bechmann, N.A., Ethridge, V.T., Gargas, N.M., Cunningham, S.D., Kuang, Z., Whicker, J.K., Rohan, J.G., Hatcher-Solis, C.N., 2022. Vagus nerve stimulation-induced cognitive enhancement: Hippocampal neuroplasticity in healthy male rats. Brain Stimul 15, 1101–1110.

Otis, J.M., Fitzgerald, M.K., Mueller, D., 2014. Infralimbic BDNF/TrkB enhancement of GluN2B currents facilitates extinction of a cocaine-conditioned place preference. J Neurosci 34, 6057–6064.

Otis, J.M., Mueller, D., 2017. Reversal of Cocaine-Associated Synaptic Plasticity in Medial Prefrontal Cortex Parallels Elimination of Memory Retrieval. Neuropsychopharmacology 42, 2000–2010.

Pena, D.F., Childs, J.E., Willett, S., Vital, A., McIntyre, C.K., Kroener, S., 2014. Vagus nerve stimulation enhances extinction of conditioned fear and modulates plasticity in the pathway from the ventromedial prefrontal cortex to the amygdala. Front Behav Neurosci 8, 327.

Peters, J., Kalivas, P.W., Quirk, G.J., 2009. Extinction circuits for fear and addiction overlap in prefrontal cortex. Learn Mem 16, 279–288.

Pitts, E.G., Taylor, J.R., Gourley, S.L., 2016. Prefrontal cortical BDNF: A regulatory key in cocaine- and food-reinforced behaviors. Neurobiol Dis 91, 326–335.

Porter, B.A., Khodaparast, N., Fayyaz, T., Cheung, R.J., Ahmed, S.S., Vrana, W.A., Rennaker, R.L., 2nd, Kilgard, M.P., 2012. Repeatedly pairing vagus nerve stimulation with a movement reorganizes primary motor cortex. Cereb Cortex 22, 2365–2374.

Reimers, J.M., Loweth, J.A., Wolf, M.E., 2014. BDNF contributes to both rapid and homeostatic alterations in AMPA receptor surface expression in nucleus accumbens medium spiny neurons. Eur J Neurosci 39, 1159–1169.

Roosevelt, R.W., Smith, D.C., Clough, R.W., Jensen, R.A., Browning, R.A., 2006. Increased extracellular concentrations of norepinephrine in cortex and hippocampus following vagus nerve stimulation in the rat. Brain Res 1119, 124–132.

Ruan, H., Yao, W.D., 2021. Loss of mGluR1-LTD following cocaine exposure accumulates Ca(2+)-permeable AMPA receptors and facilitates synaptic potentiation in the prefrontal cortex. J Neurogenet 35, 358–369.

Ruiz, A.D., Malley, K.M., Danaphongse, T.T., Ahmad, F.N., Beltran, C.M., White, M.L., Baghdadi, S., Pruitt, D.T., Rennaker, R.L., 2nd, Kilgard, M.P., Hays, S.A., 2023. Vagus Nerve Stimulation Must Occur During Tactile Rehabilitation to Enhance Somatosensory Recovery. Neuroscience 532, 79–86.

Sasase, H., Izumi, S., Deyama, S., Hinoi, E., Kaneda, K., 2019. Acute Cocaine Reduces Excitatory Synaptic Transmission in Pyramidal Neurons of the Mouse Medial Prefrontal Cortex. Biol Pharm Bull 42, 1433–1436.

Sun, W.L., Zelek-Molik, A., McGinty, J.F., 2013. Short and long access to cocaine self-administration activates tyrosine phosphatase STEP and attenuates GluN expression but differentially regulates GluA expression in the prefrontal cortex. Psychopharmacology (Berl) 229, 603–613.

Taylor, J.R., Olausson, P., Quinn, J.J., Torregrossa, M.M., 2009. Targeting extinction and reconsolidation mechanisms to combat the impact of drug cues on addiction. Neuropharmacology 56 Suppl 1, 186–195.

Tyler, R., Cacace, A., Stocking, C., Tarver, B., Engineer, N., Martin, J., Deshpande, A., Stecker, N., Pereira, M., Kilgard, M., Burress, C., Pierce, D., Rennaker, R., Vanneste, S., 2017. Vagus Nerve Stimulation Paired with Tones for the Treatment of Tinnitus: A Prospective Randomized Double-blind Controlled Pilot Study in Humans. Sci Rep 7, 11960.

Verheij, M.M., Vendruscolo, L.F., Caffino, L., Giannotti, G., Cazorla, M., Fumagalli, F., Riva, M.A., Homberg, J.R., Koob, G.F., Contet, C., 2016. Systemic Delivery of a Brain-Penetrant TrkB Antagonist Reduces Cocaine Self-Administration and Normalizes TrkB Signaling in the Nucleus Accumbens and Prefrontal Cortex. J Neurosci 36, 8149–8159.

Weiss, F., Martin-Fardon, R., Ciccocioppo, R., Kerr, T.M., Smith, D.L., Ben-Shahar, O., 2001. Enduring resistance to extinction of cocaine-seeking behavior induced by drug-related cues. Neuropsychopharmacology 25, 361–372.

Whitfield, T.W., Jr., Shi, X., Sun, W.L., McGinty, J.F., 2011. The suppressive effect of an intra-prefrontal cortical infusion of BDNF on cocaine-seeking is Trk receptor and extracellular signal-regulated protein kinase mitogen-activated protein kinase dependent. J Neurosci 31, 834–842.

Yousuf, H., Smies, C.W., Hafenbreidel, M., Tuscher, J.J., Fortress, A.M., Frick, K.M., Mueller, D., 2019. Infralimbic Estradiol Enhances Neuronal Excitability and Facilitates Extinction of Cocaine Seeking in Female Rats via a BDNF/TrkB Mechanism. Front Behav Neurosci 13, 168.

